# Isolation housing exacerbates Alzheimer’s Disease phenotype in aged APP KI mice

**DOI:** 10.1101/777524

**Authors:** M Laroy, T Saito, TC Saido, R D’Hooge, A Van der Jeugd

## Abstract

In January 2018, Britain was the first in the world to adopt a Minister of Loneliness. This illustrates the changing view on loneliness: being lonely is not just a feeling of a lack of companionship, but also a serious health problem. For example, we know that loneliness is as bad as smoking 15 cigarettes a day. Moreover, research has shown that lonely people express higher levels of cortical amyloid. Amyloid burden is an important marker of Alzheimer’s Disease (AD), a chronic neurodegenerative disease and the main cause of dementia worldwide. Together with other findings a link between loneliness, (perceived) social isolation and AD is now undeniable, but it is hard to tell from human studies whether it is the cause or the effect of AD. We need standardized animal studies to answer this question.

In an effort to study how social isolation and AD interact, we used APP KI mice bearing human transgenes known to cause AD, and isolated part of the mice in order to mimic loneliness in late-life while part of them remained group-housed. We next looked at the effects of isolation on the behaviour and symptomatology typically present in AD patients to tap cognition.

Our study reveals mixed results. Results indicate that at before isolation, at the age of 16 and 24 months, APP^NL/NL^ and APP^NL-G-F/NL-G-F^ mice do not differ to a significant extent on both the behavioural level. The APP^NL-G-F/NL-G-F^ differentiated slightly worse between the conditioned context and a new context compared to the APP^NL/NL^ mice. However, the difference appeared to be more pronounced after a period of social isolation. Social isolation had distinct effects on the AD-related anxiogenic and dementia-like phenotype. Spatial learning in the MWM task revealed distinct differences between our two models. After isolation APP^NL-G-F/NL-G-F^ mice used less spatial search strategies, compared to control mice, thus reflecting perseveration and less behavioural flexibility due to the isolation period.

## Introduction

Alzheimer’s Disease (AD) is a brain disorder with a degenerative development that makes up for the majority of all dementia cases worldwide (Alzheimer Association, 2017). Studies estimate the prevalence of dementia to be over 40 million people worldwide, with primarily people older than 60 years falling subject to the disease (Prince et al., 2013). The prevalence is projected to increase until at least 2050 by a rate of two to one every 20 years (Scheltens et al., 2016).

One of the hallmarks of AD is the presence of dense deposits that consist of *amyloid*. The deposits can be found in the functional tissue of the brain, or enclosing vascular tissue (Blennow, Mattsson, Schöll, Hansson, & Zetterberg, 2015; Crews & Masliah, 2010; Sakono & Zako, 2010). The Aβ plaques comprise redundant amounts of typical Aβ protein, which are innate to the cleavage process of the amyloid precursor protein (APP) by the β- and γ-secretase complex (Masters et al., 1985). These Aβ plaques are observed in two different states: Either in a diffuse (soluble) state or in a dense core state (Haass & Selkoe, 2007; Sakono & Zako, 2010; Selkoe, 2008; Shankar & Walsh, 2009). The toxic effects of aberrant Aβ plaques that lead to the symptoms seen in AD, start with impaired signalling between synapses (Kopeikina et al., 2011; Palop et al., 2007). Eventually, the toxicity of the aberrant Aβ plaques induces elevated abnormal electrical activity patterns all over the brain, resulting in a maladaptive neural network (Palop et al., 2007).

Social isolation (SI) or loneliness is a frequently recurring risk factor for AD, and dementia in general (Friedler, Crapser, & McCullough, 2015; Mushtaq, Shoib, Shah, & Mushtaq, 2014; Wilson et al., 2007). Social isolation is defined as “the absence or insufficient contact with others” (Friedler et al., 2015) and serves as a potential source of psychological stress. Due to its link with psychological stress, SI increases the risk for neuropsychological disorders and mortality in elderly (Friedler et al., 2015; Gilman et al., 2015; Jiang, Cowell, & Nakazawa, 2013; O’Keefe et al., 2014). The literature further suggests that SI is a potential risk factor for age-dependent cognitive impairments, dementia and more specifically AD (Leser & Wagner, 2015). One study found that SI and loneliness had a higher prevalence in AD patients compared to a group of healthy controls (El Haj et al., 2016). SI is also linked with cortical amyloid burden in healthy adults, further suggesting that SI could be a manifestation relevant to preclinical AD (Donovan et al., 2016).

The potential adverse association between SI and AD has been repeatedly investigated in rodents as well (Ali, Khalil, Elariny, & Elfotuh, 2017; Hsiao, Chen, Chen, & Gean, 2011; Hsiao, Kuo, Chen, & Gean, 2012; H. Huang et al., 2015; H. J. Huang, Liang, Ke, Chang, & Hsieh-Li, 2011). According to research, SI accelerates impairment of contextual fear memory in APP/PS1 mice by damaging long-term potentiation in CA1 neurons of the hippocampus (Hsiao et al., 2011). The same study also discovered increased levels Aβ plaques in the socially isolated mice compared to socially housed mice. In an opposite manner, group life appeared to delay the AD process in APP695/PS1-dE9 mice (Huang et al., 2015). In rats, a four-week period of SI was enough to induce neuronal deterioration and increased aggregation of Aβ plaques, as SI increased DNA fragmentation and enhanced the susceptibility for AD (Ali et al., 2017). An additional method to investigate the association between AD and SI is by looking at the structural connectome in socially isolated mice. In C57BL/6 mice, deficits in fear memory, induced by manipulation of the group size, correlated with changes in the structural connectome (Liu et al., 2016). The specific influence of SI on the progression of AD is largely unknown. SI has been shown to lead to increased production of aberrant Aβ plaques and phosphorylation of tau (Hsiao et al., 2012; Huang et al., 2015). Also, the adverse effects of social isolation are initiated by an increase in oxidative stress and an elevated inflammatory response (Powell et al., 2013). This in turn can be followed by the inhibition of anti-inflammatory responses (Azzinnari et al., 2014), synaptic plasticity (Djordjevic, Adzic, Djordjevic, & Radojcic, 2009) and myelination (Liu et al., 2012); mechanisms involved in the pathogenesis of AD.

## Methods

### Animals

Twenty-three mice, 10 APP^NL-G-F/NL-G-F^ and 13 APP^NL/NL^ mice were used. Mice originated from the Riken Institute colony (Laboratory for Proteolytic neuroscience, Riken Brain Science Institute, Japan). APP^NL/NL^ mice are considered to be a suitable negative control for the APP^NL-G-F/NL-G-F^ mice as the degree of APP should be identical in both models (Saito et al., 2014). Thus, the effects resulting from an increased Aβ42/Aβ40 ratio (Beyreuther/Iberian mutation) and increased Aβ amyloidosis (Arctic mutation) can be interpreted more straightforward. The mice were maintained under standard housing conditions, lights on at 08:00, lights off at 20:00, and provided with food and water *ad libitum*. All procedures were performed in agreement with the European Directive 2010/63/EU on the protection of animals used for scientific purposes. The protocols were approved by the Committee on Animal Care and Use at KU Leuven, Belgium (permit number: P139/2017) and all efforts were made to minimize suffering.

### Experimental Procedure

The experiment was split in two phases; the pre-isolation and isolation phase. From birth onwards, the mice were group-housed, all of the same age. Behavioural testing (Morris Water Maze and Contextual Fear Conditioning) began two weeks prior to isolation, these measurements served as the pre-isolation baseline measure of learning and memory. One day after terminating baseline testing, all mice were socially isolated for four weeks. At the end of the isolation phase, all mice were scanned at the MoSaIC Lab KU Leuven, and underwent an adapted version of the earlier conducted behavioural test battery (Reversal Morris Water Maze and Contextual Fear Conditioning).

### Behavioural tests

#### Morris Water Maze

The Morris Water Maze (MWM) test was performed prior to the isolation phase to assess hippocampal dependent spatial learning and memory (D’Hooge & De Deyn, 2001). Mice need to rely on visual spatial cues to locate a submerged platform (15 cm diameter, 5 mm beneath the surface) in a round pool (150 cm diameter) filled with opaque water (non-toxic white paint, 26 ± 1 °C). The test began with five days of acquisition training whereby each daily session existed of four trials (1-hour inter-trial interval), each trial started randomly from one of the four distinct starting locations [(day 1: 4,3,2,1), (day 2: 3,1,4,2), (day 3: 2,4,1,3), (day 4: 4,1,3,2), (day 5: 1,2,4,3)]. Mice were transported on a fly-swatter to the boundary of the maze and had to swim towards the platform, after which they were transported back to their cage underneath warming lamps. Swimming tracks were recorded via a PC-interfaced camera located above the water maze and analysed with EthoVision software (Noldus, Wageningen, The Netherlands). Mice that failed to locate the submerged platform within 120 seconds were guided via the fly-swatter towards the platform and had to remain on the platform for 10 seconds before being transported back to their cages. Two days after the last acquisition session ‘probe trials’ were conducted to assess reference memory performance. Reference memory performance is defined as the predilection for the platform area/quadrant when the platform is absent (100 seconds) (D’Hooge & De Deyn, 2001). Analyses during acquisition phase included calculating distance moved, escape latency (time-to-reach platform) and velocity via a repeated measures ANOVA (RM-ANOVA). During the probe trials analyses include time spent in each quadrant, frequency of visiting target area/quadrant and latency to reach target area/quadrant.

#### Reversal Morris Water Maze

The Reversal MWM is a shorter version of the MWM, used at the end of the isolation phase. At the start of the Reversal MWM procedure, a probe trial is conducted to assess whether there is still a remaining preference for a specific quadrant area from before the isolation phase. Then, two consecutive days with daily four trials are conducted in the same conditions as the MWM (Starting positions: 4,3,2,1; 2,4,3,1). However, the platform is placed into the opposing quadrant compared to its position during the regular MWM test earlier. At last, a final probe trial without platform (100 seconds) is conducted at the end of the second day to test reference memory performance. The same analysis procedure was used as in the regular MWM test.

#### Contextual Fear Conditioning

Context- and cue-dependent fear conditioning (CFC) is studied using an adapted protocol from Paradee et al. (1999). Electric foot shocks are applied to evoke a freezing response, which is acknowledged as a reliable measure of innate and conditioned fear in rodents (Paradee et al., 1999). The CFC test takes place in an experimental freezing set-up (Panlab, Barcelona, Spain), consisting of a test compartment (26 cm wide x 26 cm long x 27 cm high), with a grid floor placed over a startle platform and a force transducer which records animal movements. The test compartment and startle platform are located within a bigger ventilated and sound-attenuated cubicle (60 cm wide x 45 cm long x 48 cm high) with a build-in speaker. Movements of the animal are identified and recorded by the force transducer with a sampling rate of 50 Hz and saved as raw data on a PC. Two similar set-ups were used, however, to avoid confounding variables, the two groups were randomized but counterbalanced over the two set-ups. Each mouse was always tested in the same experimental set-up. The set-up is cleaned between each trial with an ethanol-based solution to ensure proper shock conductance. The pre-isolation CFC tests occurred over a period of three days. On the first day, the mice were placed in the set-up (Context A: dark, grid floor, Instanet (ethanol-based cleaning solution) and were able accommodate to the environment for five minutes during the ‘habituation phase’. During this phase, mobility is measured but no stimuli were provided. On the second day, exactly 24 hours later, the animals were placed back into the experimental set-up (Context A) and were able to acclimatize for two minutes. Subsequently, after these two minutes, two 30 second acoustic stimuli (4 kHz, 80 dB, interstimulus interval 1 minute) were presented which co-terminated with a 2 seconds 0.2 mA foot shock. This period of four minutes (2-minute acclimatization + 30 seconds stimuli and shock (1) + 1-minute interstimulus interval + 30 seconds stimuli and shock (2)) was used to calculate shock induced freezing (fear conditioning phase). Again 24 hours later, on the third day, animals were returned to the experimental set-up (Context A) for five minutes of shock-free exploration in the same context as the previous day (context test). Two hours later, the animals were returned to the test chamber in new context: Context B (lights, plastic grid cover, peppermint odour).The animals were observed for a total time of six minutes, during the first three minutes no stimulus was delivered (pre-cue score, or new context), this was followed by a three-minute presentation of the auditory cue (cued fear test). For the second assessment of the CFC after isolation, we repeated the protocol of the second and third day 24 hours apart without the habituation phase. However, for the third day we used context C (red light, soft grid cover, lavender odour) instead of context B.

## Results

### Isolation housing decreased learning and memory of aged APP KI mice

Before the SI phase, learning during the acquisition phase was observed for both groups (RM-ANOVA, Day, *F*(1,576) = 20.83, *p* < .0001) via a significant decrease in escape latency over the training days (see Figure 1A). However, the RM-ANOVA showed no statistical difference between the learning curves of the APP^NL-G-F/NL-G-F^ and APP^NL/NL^ mice during acquisition (RM-ANOVA, Genotype effect, *F*(1,576) = 0.16, *p* = .693; RM-ANOVA, “Day x Genotype” interaction, *F*(1,576) = 0.22, *p* = .639). The distance covered by the mice before escaping decreased during acquisition phase (RM-ANOVA, Day, *F*(1,576) = 59.17, *p* < .001), however no group differences were observed (RM-ANOVA, “Day x Genotype” interaction, *F*(1,576) = 1.83, *p* = .177) (see Figure 1B). Swimming speed (velocity) increased during acquisition (RM-ANOVA, day, *F*(1,576) = 74.04, *p* < .0001), again no difference between the groups was observed (RM-ANOVA, “Day x Genotype” interaction, *F*(1,576) = 3.17, *p* = .075). A probe trial was conducted two days after the last acquisition day to assess spatial reference memory. The probe trial did not show a significant preference for the target quadrant compared to the other quadrants for both APP^NL-G-F/NL-G-F^ and APP^NL/NL^ mice, on the contrary, both the APP^NL-G-F/NL-G-F^ and APP^NL/NL^ mice stayed significantly longer in the opposing quadrant, which was the starting position of the probe trial (one-way ANOVA, APP^NL-G-F/NL-G-F^, *F*(3,60) = 12.44, *p* < .0001; APP^NL/NL^, *F*(3,48) = 3.77, *p* = .016), and no group differences were observed (two-way ANOVA, “genotype x time in quadrant” interaction effect, *F*(3,108) = 1.26, *p* = .292). Their preference for the starting quadrant can be seen in the heatmaps of both groups for probe trial 1 (see Figure 2).

**Figure 1.**
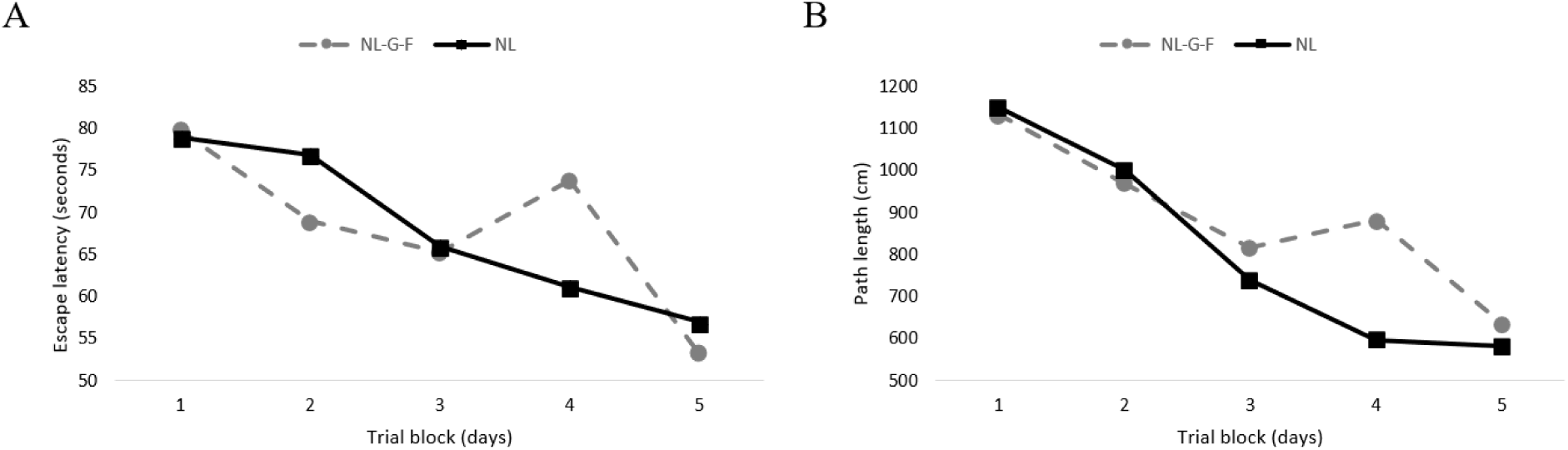
Learning curves during the acquisition phase of the MWM. **A** Time-to-reach platform decreased over the training days, however no differences are observed between the two group. **B** The covered distance during trials decreased significantly over the training days, both groups do not differ in this aspect.

**Figure 2.**
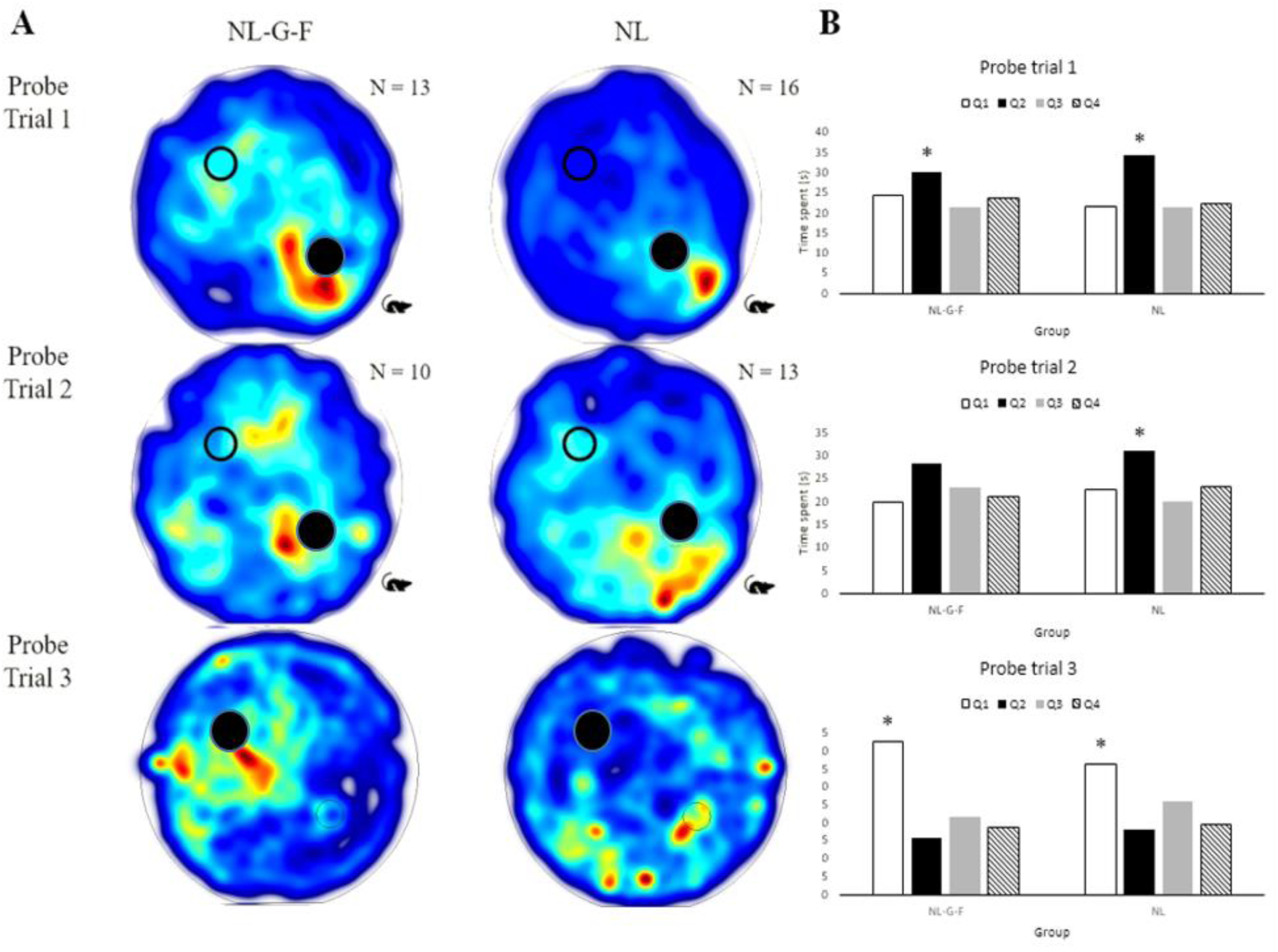
Heat maps and mean time spent in the quadrants during the probe trials in the MWM. **A** Merged group heat maps of all three probe trials, the mouse indicates the starting position, the bold black circle indicates the target area. Heat plots show insight in the dwell frequency, which is indicated by coloration from red through blue. Probe trial 1 was performed two days after the acquisition phase of the MWM. Probe trial 2 was conducted before starting with the reversal MWM and probe trial 3 was performed at the end of the reversal MWM training phase. After SI the APP^NL-G-F/NL-G-F^ mice perseverate to the previous platform position, whereas the control mice learned the new platform position. **B** Total time in the quadrants shows a clear preference for the starting quadrant (Q1 for probe trials 1 and 2, Q2 for probe trial 3) (\**p* < 0.01).

After the SI phase, the Reversal MWM protocol was used to assess spatial learning. Before the acquisition phase was performed, a probe trial (probe trial 2) was conducted to assess whether there was still a preference for a specific quadrant area. Probe trial 2 (see Figure 2) showed a significant preference of the APP^NL/NL^ mice to remain longer in the opposing quadrant, which was also the starting position of probe trial 2 (one-way ANOVA, APP^NL/NL^, *F*(3,48) = 4.66, *p* = .006). The APP^NL-G-F/NL-G-F^ showed a non-significant trend to remain longer in the starting quadrant (one-way ANOVA, APP^NL-G-F/NL-G-F^, *F*(3,36) = 2.30, *p* = .094), however no group differences were observed when comparing both groups (two-way ANOVA, “genotype x time in quadrant” interaction effect, *F*(3,84) = 0.69, *p* = .561).

Learning curves for the reversal MWM, based on escape latency over the two acquisition days, were not observed in both the APP^NL-G-F/NL-G-F^ and APP^NL/NL^ mice. Escape latency did not change significantly (RM-ANOVA, model, *F*(3,157) = 2.09, *p* = .103) during the acquisition phase, and neither did velocity (RM-ANOVA, model, *F*(3,157) = 2.60, *p* = .054). Distance travelled became significantly smaller during acquisition (RM-ANOVA, day, *F*(1,157) = 4.02, *p* = .046), however no interaction with genotype was observed (RM-ANOVA, “day x genotype” interaction, *F*(1,157) = 1.80, *p* = .182). The third probe trial (after two acquisitions days of the Reversal MWM) showed a significant preference for the target quadrant, APP^NL-G-F/NL-G-F^ showed a preference to remain in the opposing quadrant (see Figure 2A), which was the starting quadrant of the probe trial, whereas APP^NL/NL^ mice learned the new platform position (*F*(3,48) = 16.03, *p* < .0001). No group differences were observed (two-way ANOVA, “genotype x time in quadrant” interaction effect, *F*(3,84) = 1.45, *p* = .235).

### Social Isolation exacerbates fear memory of aged APP KI mice

In the CFC task before SI, both groups did not display significantly different freezing rates when exposed to the same context (context test) after the experience of a foot shock (*p* > 0.05 for all comparisons). However, the APP^NL-G-F/NL-G-F^ mice showed a higher freezing rate within the first minute of the cued fear test (exposed to a different context without cue compared to the conditioning phase) compared to the APP^NL/NL^ mice (*t*(27) = −2.06, *p* = .049) (see Figure 3A). From the second to sixth minute of the cued fear test no differences were observed between the two groups (*p* > 0.05 for all comparisons). When looking at the average freezing rates in the two phases of the cued fear test (no auditory cue: First to third minute; with auditory cue: Fourth to sixth minute) no significant differences are observed between the two groups.

**Figure 3.**
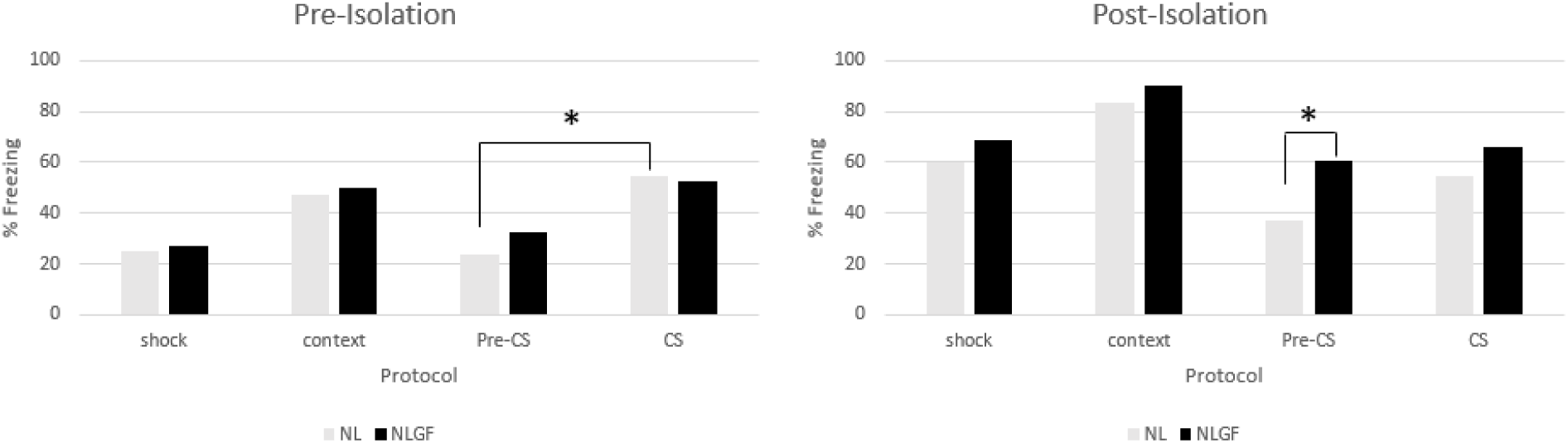
Mean freezing rates (%) during the CFC task. Before SI, the APP^NL/NL^ mice displayed a significant different between the pre-conditional and conditional stimulus phase in the new context. After SI, a significant difference is observed in freezing between the APP^NL/NL^ and APP^NL-G-F/NL-G-F^ mice during the pre-conditional stimulus phase in the new context (\**p* < 0.01).

After SI, groups did not display a significant difference in freezing rates when exposed to same context after the experience of a foot shock (context test). However, when introduced to a new context (cued fear test), the APP^NL-G-F/NL-G-F^ mice showed a higher freezing rate within the first and second minute (without auditory cue) (first minute, *t*(21) = −2.57, *p* = .018; second minute, *t*(21) = -3.59, *p* = .002). When looking at the average freezing rates in the two phases of the cued fear test (no auditory cue: First to third minute; with auditory cue: Fourth to sixth minute), the APP^NL-G-F/NL-G-F^ mice showed a significant higher freezing rate during the first phase (new context without auditory cue) compared to the APP^NL/NL^ mice (*t*(21) = −2.97, *p* = .007) (see Figure 3B). No significant differences are observed in the second phase (new context with auditory cue).

## Discussion

A new strain of knock-in (KI) mice (APP^NL-G-F/NL-G-F^) carrying Swedisch (NL), Beyreuther/Iberian (F), and Arctic (G) mutations in the APP gene are used for our study. Each mutation has its own corresponding pathological feature. The Swedish mutation promotes expression of Aβ40 and Aβ42 and the Beyreuther/Iberian mutation elevates the ratio of Aβ42 to Aβ42, which further increases the expression of Aβ42. The newly added Arctic mutation stimulates Aβ amyloidosis limited to areas with neuroinflammatory responses (Masuda et al., 2016; Saito et al., 2014). Hence, APP^NL-G-F/NL-G-F^ mice produce APP similar to APP^NL-F/NL-F^ mice, resulting in increased Aβ40 and Aβ42 expression and an increased Aβ42 to Aβ40 ratio in an age-dependent manner, assuming that there are no interactions with the additional Arctic mutation. The additional value of the APP^NL-G-F/NL-G-F^ model originates from the Arctic mutation. This mutation causes destructive Aβ amyloidosis by altering the binding process of antibodies to Aβ (Saito et al., 2014). The APP^NL/NL^ model with only the Swedish mutation is frequently used as a control model since the Aβ deposition and amyloidosis are insignificant (Saito et al., 2014). While Saito et al. (2014) demonstrated the increased value of the new APP KI model for experimental studies of AD, characterization of the behavioural phenotype of the model remains inadequate. Concluding that the APP^NL-G-F/NL-G-F^ mice showed typical Aβ pathology, neuroinflammation and memory impairment in previous research (Masuda et al., 2016; Saito et al., 2014), this strain can serve as an exemplary medium for studying pathological mechanisms of AD.

Subtle effects were found between the two strains before SI. CFC revealed a worse performance for the APP^NL-G-F/NL-G-F^ when placed in a new context, however the difference appeared to be more pronounced after a period of SI. Spatial learning in the MWM task did not reveal any differences between our two strains. Before SI, both strains showed a learning curve in the MWM during acquisition. Over five days of training, the escape latency decreased significantly in both groups. It is surprising that 16- and 23 months old APP KI mice were still able to learn to some extent, considering the extent of their predicted Aβ pathology (Masuda et al., 2016; Saito et al., 2014). However, the probe trials revealed that the learning effects during acquisition did not translate to a preference for the target quadrant two days after the last training day. During the probe trials, both groups spent most of the time in the starting quadrant (opposing quadrant). Both groups could not recall the place of the platform, however there was a statistical difference in the frequency of visits of the target quadrant, meaning that the APP^NL-G-F/NL-G-F^ mice visited the target quadrant more often compared to the APP^NL/NL^ mice. However, the time spent in the target quadrant does not differ between the groups, hence we cannot infer that the APP^NL-G-F/NL-G-F^ mice recall the place of the platform better than the APP^NL/NL^ mice.

However, after SI, only the control group showed clear signs of spatial learning via the reversal MWM. The absence of learning effects could be caused by our relatively small sample size of 23 mice (10 APP^NL-G-F/NL-G-F^ mice and 13 APP^NL/NL^ mice). It could be argued that aging effects are causing the deficiency in learning effects. Before SI, both the APP^NL-G-F/NL-G-F^ and APP^NL/NL^ mice displayed the same freezing rates during the context test (same context as during conditioning phase), therefore we can conclude that both groups learned the association of the context and the foot shock. Group differences were observed during the cued fear test. Within the first minute in a new context without the auditory cue, the APP^NL-G-F/NL-G-F^ mice showed increased freezing rates compared to the APP^NL/NL^ mice. The APP^NL-G-F/NL-G-F^ mice cannot make the distinction between the new and old context, the APP^NL/NL^ mice do seem to be able to make the distinction between the context from the fear conditioning phase and the new context. However, when the auditory cue is introduced within the new context, both groups show the same freezing rates. For the APP^NL-G-F/NL-G-F^ mice, the amount of freezing during the cued fear test is the same with and without the auditory cue. For the APP^NL/NL^ mice, the freezing rates between the cued fear test with and without auditory cue are significantly different. Hence, the APP^NL/NL^ mice seem to make the distinction between the original context and new context in the absence of the auditory cue, a distinction the APP^NL-G-F/NL-G-F^ mice do not seem to make.

The CFC task was repeated after a period of SI. Both groups displayed the same freezing rates in the context test, both strains do seem to learn the meaning of the context, which is associated with a pain stimulus. However, the difference between the APP^NL-G-F/NL-G-F^ and APP^NL/NL^ mice looks to be less pronounced during the cued fear test. The amount of freezing during the cued fear test without auditory cue compared the presence of the auditory cue is no longer significantly different for both groups. The APP^NL-G-F/NL-G-F^ mice have the same freezing rates in the two blocks. For the APP^NL/NL^ mice there appears to be a trend for a difference in freezing rates with and without auditory cue, but this is not significant.

Indeed, the main psychological symptom observed in AD is indeed spatial memory impairment (like in MWM test), but now we speculate that recall may be influenced when an emotional component is associated with a (negative) event. In AD subjects this can even be enhanced through social isolation, as our (single) foot-shock experiment show. According to our findings, isolated housing is a risk factor for memory dysfunctions. It can be concluded that results concerning the effects of SI on memory are not equivocal. While long-term spatial memory is unaffected by SI, there seems to be an impact on long-term fear memory. This needs further research into the mechanisms of memory formation in healthy and disease, and in loneliness as well as ‘normal’ social settings.

We can conclude, that if social isolation is a risk factor for AD, it therefore could be a modifiable one. Targeting isolated individuals of all ages may therefore have the potential to decrease AD in the population, which is currently one of the ultimate priorities of our global public health. Also the possibility that re-socialization after a period SI is able to undo any negative effects demands further investigation. If SI has a deleterious impact on cognition, re-socialization, physical or cognitive stimulation could have the potential of functioning as a protective treatment of AD.

